# Identification of zinc-dependent mechanisms used by Group B *Streptococcus* to overcome calprotectin-mediated stress

**DOI:** 10.1101/2020.08.14.252064

**Authors:** Lindsey R. Burcham, Yoann Le Breton, Jana N. Radin, Brady L. Spencer, Liwen Deng, Aurélia Hiron, Monica R. Ransom, Jéssica da C. Mendonça, Ashton T. Belew, Najib M. El-Sayed, Kevin S. McIver, Thomas E. Kehl-Fie, Kelly S. Doran

**Affiliations:** Department of Immunology and Microbiology, University of Colorado School of Medicine, Aurora, CO, USA; Cell Biology and Molecular Genetics, University of Maryland, College Park, MD; Department of Microbiology, University of Illinois at Urbana-Champaign, Urbana, IL, USA; Carl R. Woese Institute for Genomic Biology, University of Illinois at Urbana-Champaign, Urbana, IL, USA; Université de Tours, INRAE, ISP, F-37000, Tours, France; Current address: Walter Reed Army Institute of Research, Silver Spring, MD, USA; Center for Bioinformatics and Computational Biology, University of Maryland, College Park, MD, USA

## Abstract

Nutritional immunity is an elegant host mechanism used to starve invading pathogens of necessary nutrient metals. Calprotectin, a metal binding protein, is produced abundantly by neutrophils and is found in high concentrations within inflammatory sites during infection. Group B *Streptococcus* (GBS) colonizes the gastrointestinal and female reproductive tracts and is commonly associated with severe invasive infections in newborns such as pneumonia, sepsis, and meningitis. Though GBS infections induce robust neutrophil recruitment and inflammation, the dynamics of GBS and calprotectin interactions remain unknown. Here we demonstrate that disease and colonizing isolate strains exhibit susceptibility to metal starvation by calprotectin. We constructed a *mariner* transposon (*Krmit*) mutant library in GBS and identified 258 genes that contribute to surviving calprotectin stress. Nearly 20% of all underrepresented mutants following treatment with calprotectin, are predicted metal transporters, including known zinc systems. As calprotectin binds zinc with picomolar affinity, we investigated the contribution of GBS zinc uptake to overcoming calprotectin-imposed starvation. Quantitative RT-PCR revealed a significant upregulation of genes encoding zinc-binding proteins, *adcA*, *adcAII*, and lmb, following calprotectin exposure, while growth in calprotectin revealed a significant defect for a global zinc acquisition mutant (Δ*adcA*Δ*adcAII*Δ*lmb*) compared to the GBS WT strain. Further, mice challenged with the Δ*adcA*Δ*adcAII*Δ*lmb* mutant exhibited decreased mortality and significantly reduced bacterial burden in the brain compared to mice infected with WT GBS; this difference was abrogated in calprotectin knockout mice. Collectively, these data suggest that GBS zinc transport machinery are important for combatting zinc-chelation by calprotectin and establishing invasive disease.

**Importance:** GBS asymptomatically colonizes the female reproductive tract but is a common causative agent of meningitis. GBS meningitis is characterized by extensive infiltration of neutrophils, carrying high concentrations of calprotectin, a metal chelator. To persist within inflammatory sites and cause invasive disease, GBS must circumvent host starvation attempts. Here, we identified global requirements for GBS survival during calprotectin challenge, including known and putative systems involved in metal ion transport. We characterized the role of zinc import in tolerating calprotectin stress *in vitro*, and in a mouse model of infection. We observed that a global zinc-uptake mutant was less virulent compared to the parental GBS strain and found calprotectin knockout mice to be equally susceptible to infection by WT and mutant strains. These findings suggest that calprotectin production at the site of infection results in a zinc-limited environment and reveals the importance of GBS metal homeostasis to invasive disease.

## Introduction

Bacteria, like eukaryotes, have a strict requirement for transition metals that often function as enzyme cofactors or provide protein structural support (1). Though essential for survival, metal ions can also be toxic, and to successfully survive within a host, pathogens must coordinate ion uptake and efflux to maintain intracellular metal homeostasis (2–4). To antagonize the nutritional requirements of invading pathogens, the vertebrate host immune system has evolved elaborate mechanisms for restricting access to metal ions, a process termed nutritional immunity (5, 6). The hosts’ efforts to limit access to metal ions can dampen pathogen metalloenzyme function, restricting growth and the ability to cause disease (7–9). Widely recognized host iron-binding proteins include transferrin, lactoferrin, and lipocalin-2 that sequester iron(III) or iron-bound siderophores (10–12) from pathogens. Calprotectin, another metal-binding host protein is unique in that it can interact with multiple metal ions (2). Calprotectin is a tetraheterodimer of two members of the S100 protein family, S100A8/S100A9, or calgranulin A/B and MRP-8/14 (6, 13), and makes up approximately 50% of the neutrophilic cytoplasmic protein content (13). The S100A8 and S100A9 form a heterodimer that upon calcium-dependent conformational change, create two metal-binding sites that bind zinc with picomolar/femtomolar affinity (14–16) and manganese at nanomolar affinities (7, 16, 17). More recent studies have shown that calprotectin can additionally chelate iron(II) (18–20), copper (20, 21), and nickel *in vitro* (22) but the implications of the binding of these metals during infection is not understood. Calprotectin is abundant during inflammation or at sites of infection, where concentrations can exceed 1 mg/mL; therefore, invading pathogens must be able to cope with these pressures to cause disease (17, 23).

*Streptococcus agalactiae*, or Group B *Streptococcus* (GBS), is a pathobiont that colonizes the vaginal tract but can be a severe threat to the fetus and newborn. The onset of GBS invasive disease in the neonate can occur as a result of aspiration during passage through a colonized birth canal (24), bacterial transmigration through the bloodstream (25), and penetration of the blood-brain barrier (BBB) (26). To combat the risk of infection in newborns, many countries have implemented the use of prophylactic antibiotics administered to colonized pregnant mothers at the time of delivery (27); however, despite these widespread efforts, GBS remains a leading cause of neonatal pneumonia, sepsis, and meningitis (28). Bacterial meningitis is a severe and potentially lethal pathology of the central nervous system that develops when pathogens overcome host defenses and successfully penetrate the BBB. Meningitis is characterized by an overwhelming cytokine response and immune cell influx to the site of infection (29). Meningitis is a particularly complex disease and results in a neonatal mortality rate as high as 40% (29, 30). Further, the associated inflammation results in neuronal damage and brain injury, with nearly 20-50% of surviving patients suffering permanent neurological sequelae including hearing and vision impairment, cognitive deficiencies, and seizures (31, 32).

During acute bacterial meningitis, neutrophils predominate in the cerebral spinal fluid, which is often used as a diagnostic marker. Following interaction with human cerebral microvascular endothelial cells (hCMEC) *in vitro,* GBS induces a characteristic neutrophilic inflammatory response, including expression of chemoattractants interleukin-8 (IL-8), C-X-C motif chemokine ligand 1 (CXCL-1), and CXCL-2 (33–35). Similar results are observed in animal models of experimental GBS meningitis, as brain tissue of GBS infected mice shows increased neutrophil and monocyte infiltration compared to naïve controls (35) indicating a close interaction between GBS and granulocytic cells during active infection. Additional studies have shown that, in response to GBS, neutrophils elaborate extracellular traps decorated with lactoferrin (36) and that S100A9, a calprotectin subunit, is present in the blood and amniotic fluid during intrauterine GBS infection (37). These observations suggest that GBS experiences metal limitation during infection, but the mechanisms used by GBS to overcome nutritional immunity remain unknown.

Bacteria utilize a number of strategies to obtain zinc during infection including direct uptake of the metal, the use of metallophores, and piracy from zinc-bound host proteins. While there are a myriad of strategies employed to obtain zinc, the AdcABC/ZnuABC family of ATP-binding cassette transporters are present in most bacteria. Streptococcal pathogens *S. pneumoniae* and *S. pyogenes* encode two zinc-binding proteins AdcA and AdcAII, whereas GBS is particularly distinct as it possesses a third zinc-binding protein, Lmb (38, 39). AdcA and AdcAII/Lmb have been shown to utilize distinct mechanisms to bind zinc ions and shuttle them through the AdcBC transporter and are important for growth in zinc-restricted environments and infection (40–43).

Here we investigated GBS fitness during calprotectin stress using a newly constructed saturated transposon mutant library and targeted amplicon sequencing. We characterized the global requirements for GBS survival during nutritional immunity, identifying 258 mutants, 123 underrepresented and 135 overrepresented, that impact calprotectin sensitivity. We show herein that characterized and putative metal transporters are important for calprotectin survival *in vitro*, and that the zinc uptake machinery contributes to survival during calprotectin-induced starvation and invasive disease progression. These results provide insight into zinc-dependent mechanisms that GBS employs to evade the host immune response and nutritional immunity to successfully cause disease, and establish a groundwork to study the comprehensive effects of chelation on multi-metal transport in GBS.

## Results

### GBS growth inhibited in the presence of calprotectin

Previous studies have shown calprotectin inhibits bacterial growth by limiting nutrient metal ions (7–9). To characterize the response of GBS to metal chelation by calprotectin, we assessed growth of GBS strains in the presence of purified calprotectin. Disease clinical isolates A909 (serotype Ia) (44), CJB111 (serotype V) (45), and COH1 (serotype III) (46), were incubated with increasing concentrations of purified calprotectin ranging from 0-480 μg/mL and growth was assessed by optical density and plating for viable bacteria. Though varying levels of chelation sensitivity were detected, growth of all GBS strains (as measured by OD_600nm_) was significantly inhibited at high, but still physiologically relevant, calprotectin doses (Fig 1A-C). Similar patterns of growth inhibition were observed when growth was assessed by enumerating CFU following an 8-hour incubation with calprotectin. Exposure to calprotectin at concentrations higher than 120 μg/mL significantly inhibited growth of all GBS strains (Fig 1D-F). Additionally, we determined how a panel of 27 vaginal isolates collected from the vaginal tracts of pregnant women (47) survived in the presence of calprotectin. All strains were grown with or without 120 μg/mL calprotectin, and data are displayed as percent inhibition in treated versus untreated controls for isolates across five different capsular serotypes. We observed a mean percent inhibition across the isolates that ranged from 30-55%. When mean inhibition of each serotype was compared against the others, we observed that vaginal isolates belonging to serotype V exhibited the most variability between strains and were significantly more resistant to calprotectin-mediated chelation than serotype Ia isolates (Fig 1G). Representative invasive isolates A909, CJB111, and COH1 were included in the isolate panel in their capsular serotype grouping and are denoted by the open shape. These data suggest that while there is variation in levels of sensitivity across strains and serotypes, GBS is broadly sensitive to the antimicrobial activity of calprotectin.

**Figure 1.**
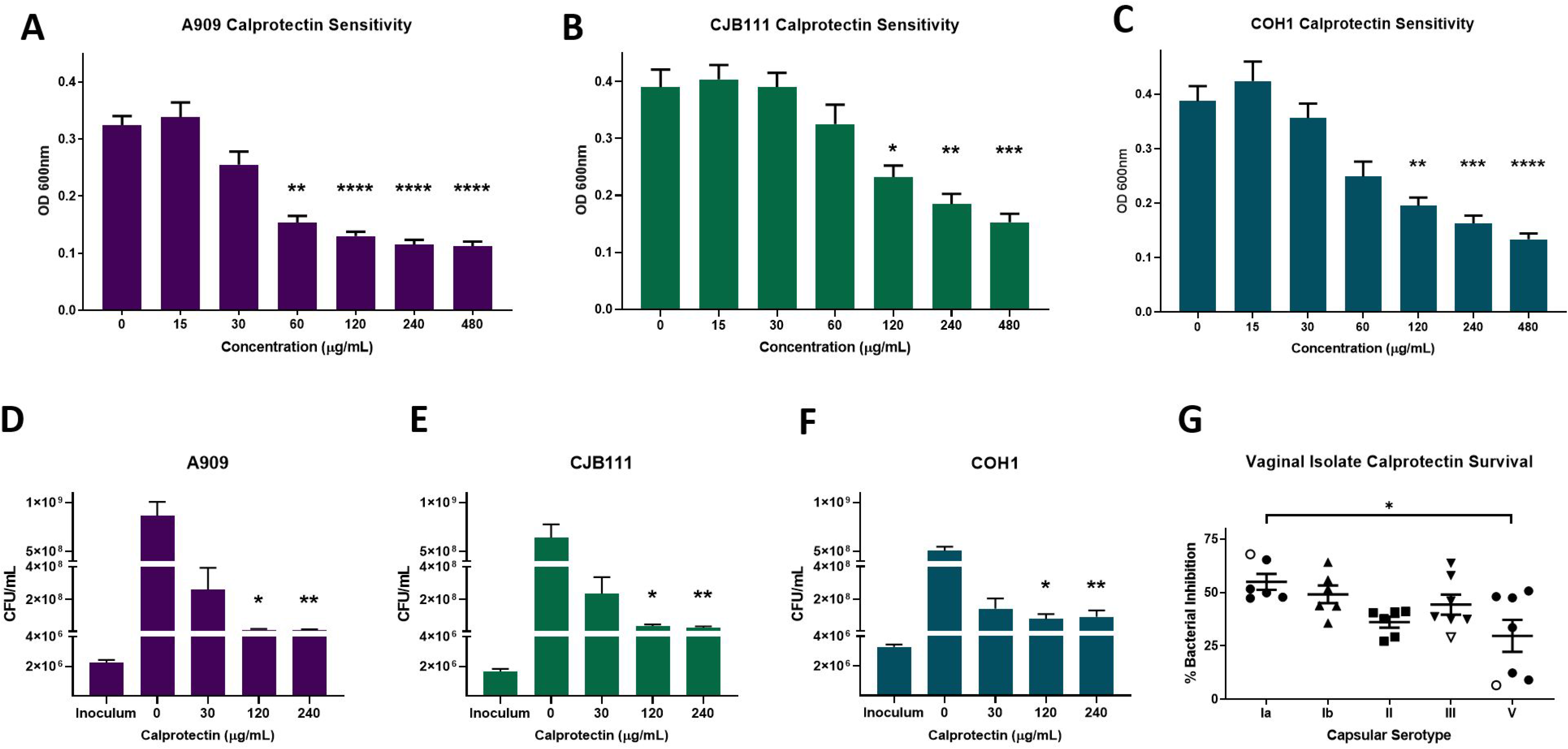
Calprotectin inhibits GBS growth *in vitro*. Growth of GBS invasive isolates **A)** A909, **B)** CJB111, and **C)** COH1 were assessed by measuring optical density (OD_600nm_) following 8-hour incubation with recombinant calprotectin (0-480 μg/mL) or by quantitating CFU **(D-F)**. **G)** Sensitivity was assessed by OD_600nm_ across a panel of vaginal isolates (closed shapes) and invasive isolates (open shapes) following an 8-hour incubation with 120 μg/mL calprotectin. Data are displayed as percent growth inhibition as compared to untreated isolate controls. All experiments were performed in technical triplicates of *n*=3 and data were averaged together from three independent experiments. Significance for panels A-F was determined by Kruskal-Wallis with Dunn’s multiple comparisons test comparing treated samples to untreated controls. Significance for panel G was determined by One-way ANOVA with Tukey’s multiple comparisons test, * *p*<0.05, ** *p*<0.01, *** *p*<0.001, **** *p*<0.0001.

### Essential genes for GBS growth in calprotectin

In order to successfully colonize the host or survive within highly inflammatory environments during infection, GBS must cope with nutritional immunity and specifically metal-limitation imposed by calprotectin. To identify factors that are important for responding to calprotectin-mediated chelation, we constructed a GBS saturated *Krmit*-Tn mutant library in the CJB111 strain background as described previously for Group A *Streptococcus* (48). Analysis of the library revealed 68,857 unique insertion sites across the GBS genome (Fig 2). To identify essential genes for GBS growth, we outgrew the Tn-mutant library in Todd Hewitt Broth with yeast extract (THY), modified-RPMI (mRPMI), and mRPMI plus subinhibitory (60 μg/mL) and inhibitory (480 μg/mL) concentrations of calprotectin. We recovered CFU from these growth conditions, extracted genomic DNA, and prepared sequencing libraries as described in the methods. Transposon insertions were sequenced as previously described (49) with minor changes, and sequenced reads were mapped back to the GBS genome. Bayesian statistical analyses (50) identified in the absence of calprotectin, 206 essential genes for growth in THY and 450 essential genes for growth in mRPMI (Fig 3A, B), with 153 essential genes common to both media conditions (Fig 3C, Supplemental Table 1). The genes deemed essential for growth in THY and mRPMI were assigned clusters of orthologous groups of proteins (COGs) and were found to be involved primarily in translation, ribosomal structure, and biogenesis (23% in THY, 13% in mRPMI); replication, recombination, and repair (16% in THY, 13% in mRPMI); cell wall/membrane/envelope biogenesis (10% in THY, 10% in mRPMI); and carbohydrate transport and metabolism (10% in THY, 7% in mRPMI) (Fig 3D, E).

**Figure 2.**
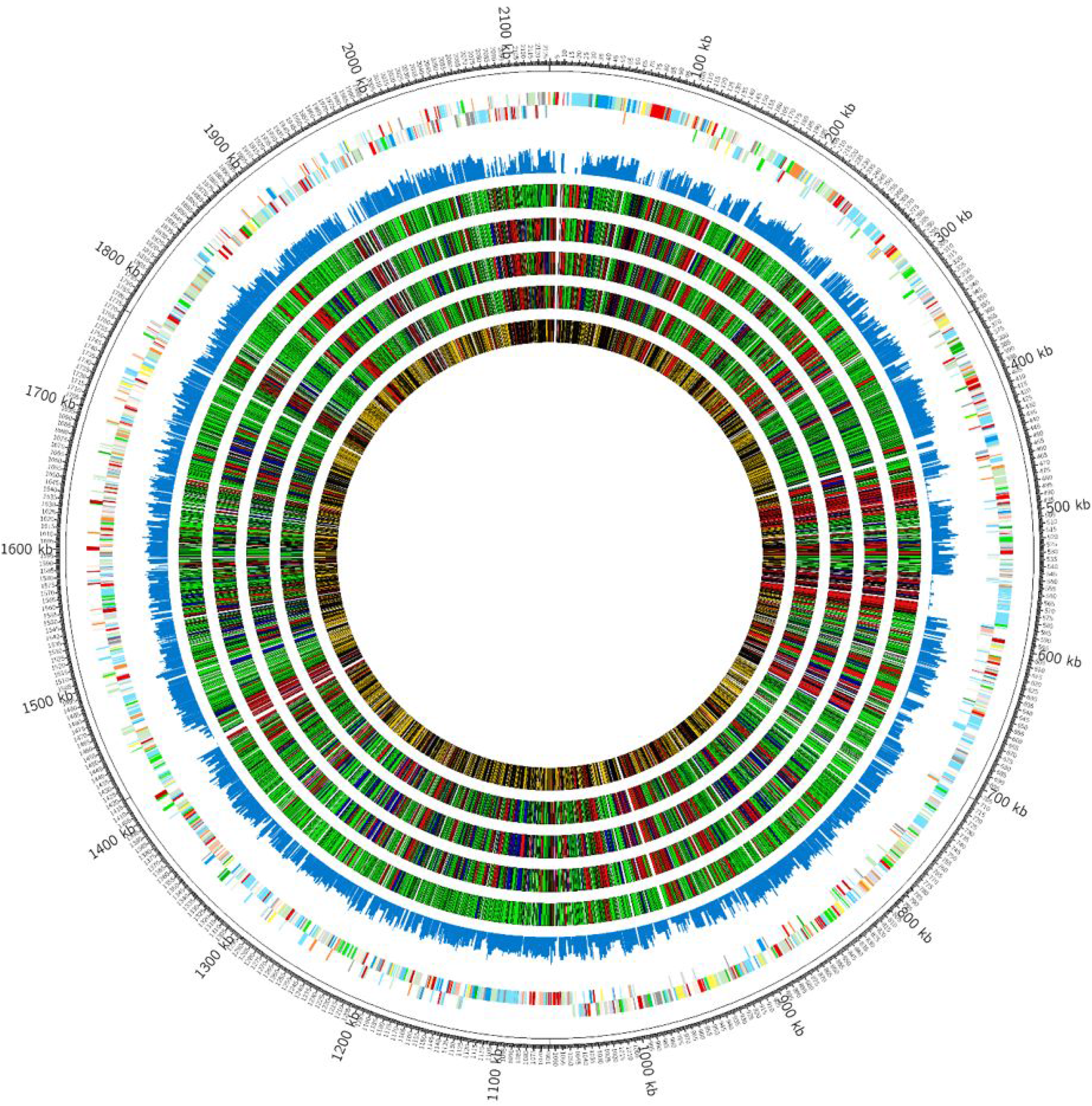
Construction of a saturated *Krmit*-transposon mutant library in GBS. CIRCOS atlas representation of the A909 genome is shown with base pair (bp) ruler on outer ring. The next two interior circles represent GBS open reading frames on the (+) and (−) strands, with colors depicting COG categories. The next circle (blue) indicates the frequency of Krmit TIS observed in the initial mutant library grown in THY, with 68,857 unique insertion sites detected. The inner four circles present the results of Bayesian analysis of GBS gene essentiality in different growth conditions, (THY, mRPMI, mRPMI plus subinhibitory calprotectin, mRPMI plus inhibitory dose calprotectin in order towards center; essential genes (red), non-essential genes (green), and excluded genes in either grey (too small for analysis) or black (inconclusive call). The center circle compiles the summary analysis of GBS genes in all four growth conditions, with essential genes in all conditions (red), non-essential genes in all conditions (yellow) and small/inconclusive genes (grey/black).

**Figure 3.**
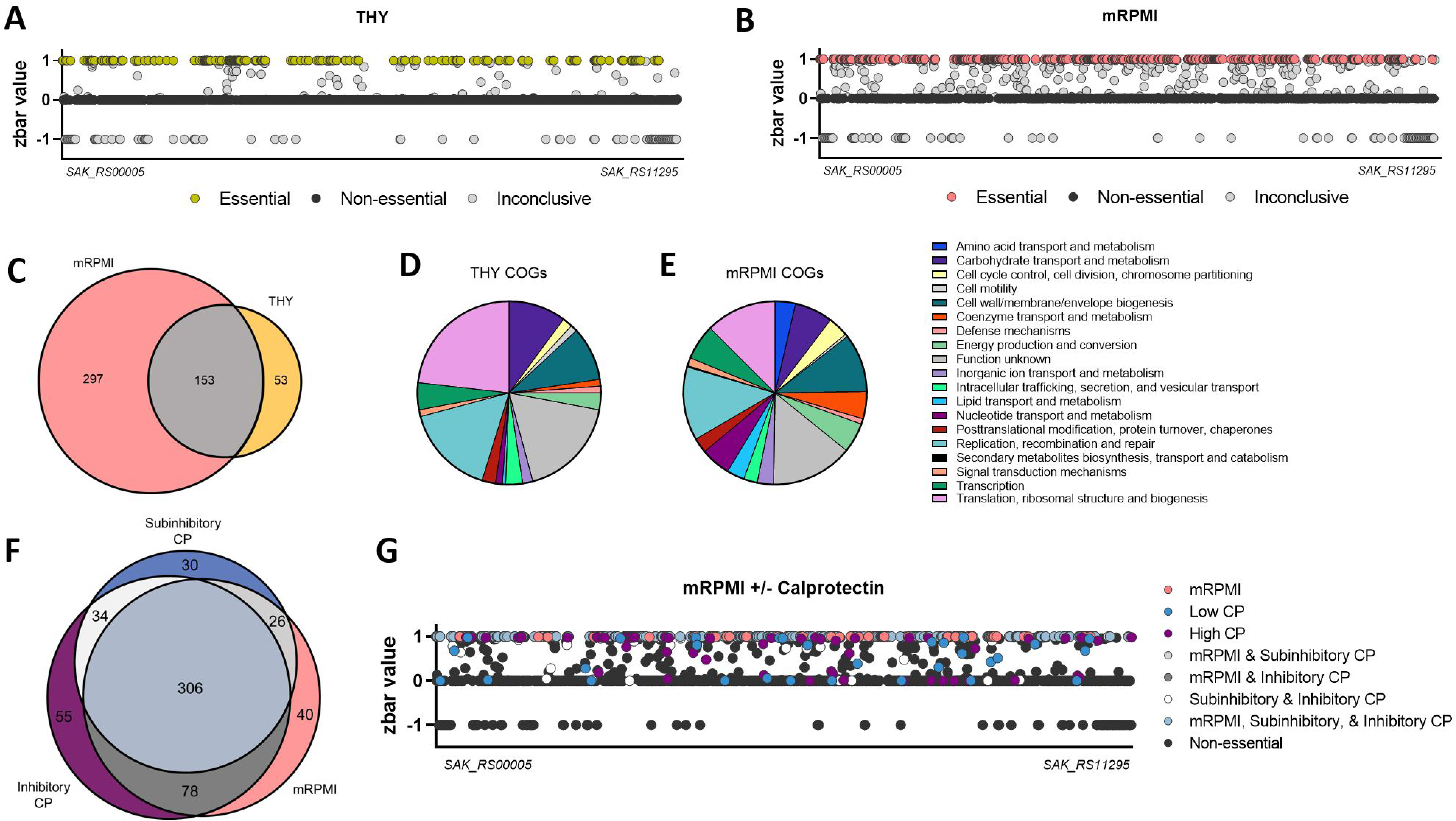
GBS essential genes for growth *in vitro*. Bayesian analysis of essential genes for growth in **A)** THY or **B)** mRPMI. Essential genes are depicted as **A)** yellow or **B)** pink, non-essential genes are shown in black, and inconclusive genes are shown in gray. The X-axis is a linear representation of the A909 genome. EggNOG 5.0 was used to assign COGs to determine functions for essential genes for growth in **D)** THY and **E)** mRPMI. Venn diagrams depict the essential genes for growth in **C)** mRPMI and THY or **F)** mRPMI, subinhibitory (60 μg/mL), and inhibitory (480 μg/mL) calprotectin. **G)** Linear map represents Bayesian analyses of essential genes for growth in mRPMI, subinhibitory and inhibitory calprotectin.

Bayesian analyses were then used to determine the essential genes for growth in the presence of calprotectin. These analyses compared essential genes from the base media (mRPMI) and subinhibitory and inhibitory concentrations of calprotectin. From these analyses, we identified 40 genes that were essential specifically for growth in mRPMI, 30 genes that were essential only for growth in a subinhibitory dose of calprotectin, and 55 genes that were essential only for growth in an inhibitory calprotectin dose. The remaining 306 essential genes were deemed important for growth across all environments (mRPMI, subinhibitory calprotectin, and inhibitory calprotectin) (Fig 3F, 3G, Supplemental Table 1).

### Global impact of calprotectin on GBS fitness

To determine the global effect of calprotectin on GBS fitness, differential analyses were performed using DESeq2 comparing subinhibitory (60 μg/mL) or inhibitory (480 μg/mL) calprotectin treated samples to untreated mRPMI controls. Genes found to be essential for growth in mRPMI (pink), subinhibitory calprotectin (blue), inhibitory calprotectin (purple), or across two treatments (gray) were excluded from fitness analyses (Fig 4A, B). We characterized the global impact of calprotectin stress on GBS growth and identified a total of 258 mutants in the output pool whose growth was significantly impacted by calprotectin. We identified 135 mutations that conferred a fitness advantage for GBS during calprotectin stress, with 94 mutants in subinhibitory dose and 98 mutants in inhibitory dose calprotectin (Fig 4A, B, Table S1), with 57 mutants common between the two calprotectin treatment groups. We also identified 123 mutants that resulted in a fitness defect during calprotectin stress, and of those, 93 mutants were important in subinhibitory dose of calprotectin and 76 mutants were important for in inhibitory levels of calprotectin (Fig 4A, B, Table S1), with 46 mutants observed in both calprotectin-treated samples (Fig 4C). Of the Tn-mutants that were identified as underrepresented following treatment with calprotectin, COGs were identified for 76 of the mutations observed in subinhibitory treatment and 60 of the mutations observed in inhibitory calprotectin treatment. The most abundant COGs of known function were those involved in inorganic ion transport, amino acid and carbohydrate transport and metabolism, and defense (Fig 4D, E). Approximately 15% of the mutants underrepresented in both concentrations were grouped into the inorganic ion COG and were previously characterized or putative systems involved in metal ion uptake or efflux (Table 1). These systems involved in maintaining metal homeostasis were many of the most significantly underrepresented mutants as denoted in the volcano plots (shown in red) for each calprotectin concentration (Fig 4A, B).

**Figure 4.**
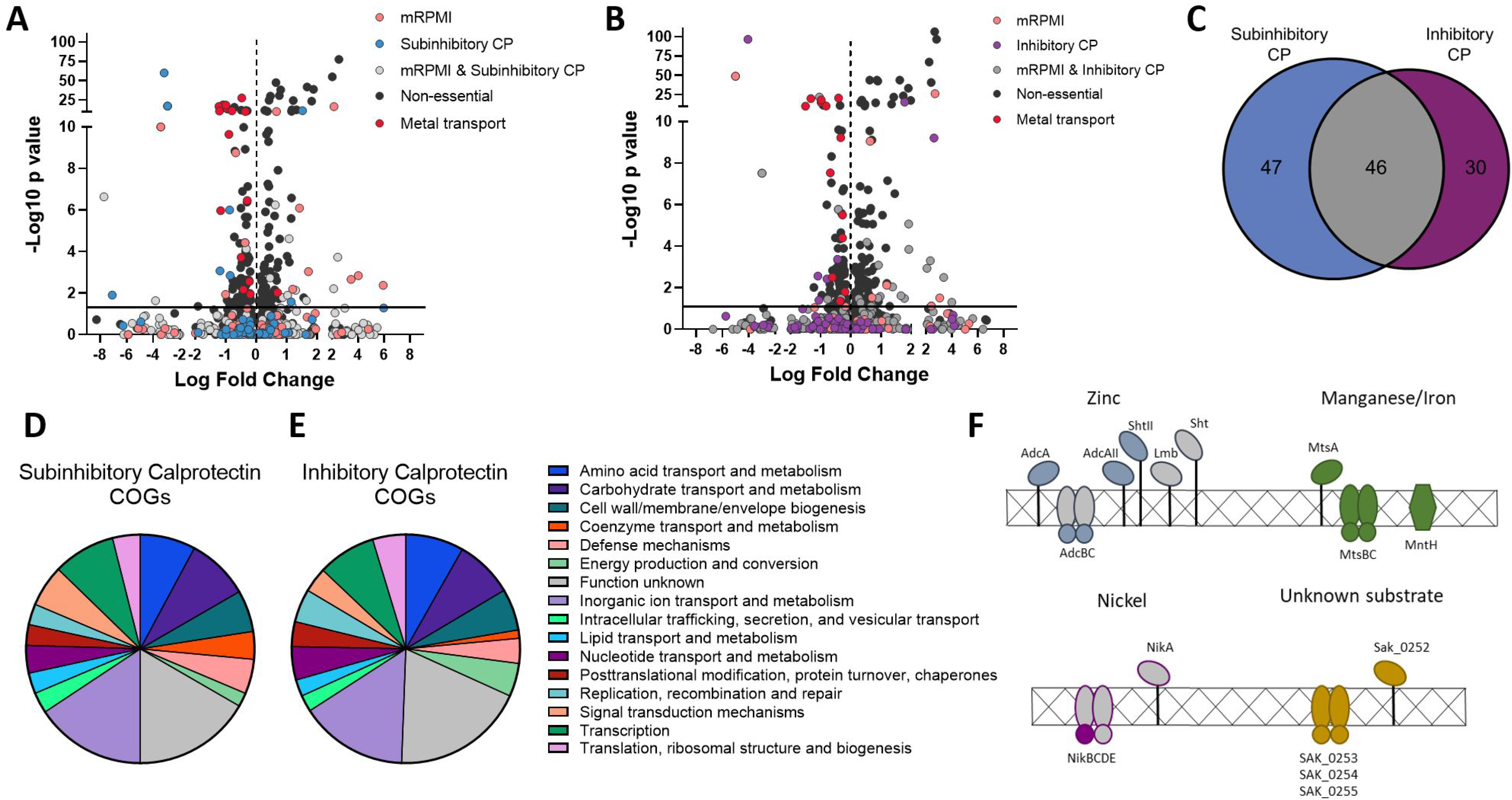
GBS genomic fitness screen in calprotectin. Volcano plots identify essential genes for growth in mRPMI and **A)** subinhibitory (60 μg/mL) calprotectin and **B)** inhibitory (480 μg/mL) calprotectin (pink, blue/purple, gray), non-essential genes (black), and genes involved in metal transport (red). (37). Essential genes (all but “nonessential” and “metal transport” were excluded from fitness analyses by DESeq2. **C)** Venn diagram depicts the underrepresented mutants detected in low (60 μg/mL) and high calprotectin (480 μg/mL) and common genes important for growth in both conditions. COGs were assigned to genes that contribute to survival in **B)** subinhibitory dose and **C)** inhibitory dose calprotectin using EggNOG 5.0. **F)** Schematic of GBS metal importers that contribute to GBS survival during calprotectin stress.

To gain more insight into how the identified metal transport systems may contribute to homeostasis, all proteins of interest were clustered by ortholog against characterized metal transport machinery in the closely related *Streptococcus pneumoniae* TIGR4 genome. Of the three genes encoding zinc-binding proteins, we identified only *adcA* (*SAK_0685*) and *adcAII* (*SAK_1898*) as important for growth in calprotectin. The gene encoding their cognate ATPase, *adcC* (*SAK_0218*) and one of the genes encoding a streptococcal histidine triad protein, *shtII* (*SAK_1897*), were also underrepresented in our Tn-sequencing analyses (Fig 4F). Underrepresentation of mutants in *adcAII* was specific to treatment with inhibitory levels of calprotectin, while mutants in the remaining zinc transport genes detected were defective in calprotectin survival independent of concentration (Table 1).

In addition to the zinc machinery, significant underrepresentation was detected for Tn-insertions in the genes encoding the manganese/iron ABC-transporter, *mtsABC* (*SAK_1554-1556*), and the gene encoding the manganese/iron natural resistance-associated macrophage protein (NRAMP), *mntH* (*SAK_0871*) (51) (Fig 4F). Ortholog clustering again confirmed the conservation of the GBS transporter MtsABC to the pneumococcal transporter PsaABC (51, 52); however, other pathogenic streptococci including *S. pyogenes* and *S. pneumoniae* are devoid of manganese- and iron-dependent NRAMP transporters (53–55). Additionally, mutants in genes that comprise two other putative metal ion uptake systems were identified as underrepresented (Fig 4F). *nikD* (*SAK_1539*) is an ATP-binding protein encoded within the operon *nikABCDE* (*SAK_1538-1542*) that encodes a putative but uncharacterized nickel ABC transport system, and the second underrepresented and uncharacterized ABC-transport system is encoded by *SAK_0252-0255*; however, the substrate transported by this system remains unknown (Fig 4F). The substrate binding proteins of both putative metal transport systems belong to the NikA/DppA/OppA superfamily.

### Calprotectin induces expression of zinc import machinery

Upon identifying mutants in genes encoding two zinc-binding proteins, *adcA* and *adcAII*, in our transposon sequencing analyses, we hypothesized that differential expression of genes involved in zinc acquisition might occur following exposure to calprotectin, a natural source of zinc limitation. Quantitative RT-PCR analysis of genes encoding the zinc-binding proteins *adcA*, *adcAII*, and *lmb*, was performed following treatment with calprotectin or the cell membrane permeable chelator N,N,N’,N’-tetrakis(2-pyridinlymethyl)-1,2-ethanediamine, (TPEN). Fold changes in gene expression were calculated by ΔΔCT with *gyrA* serving as an internal control and were compared to expression observed in untreated controls. Expression of *adcA* was induced by 4-fold following exposure to calprotectin and 10-fold following TPEN treatment, while expression of *adcAII* and *lmb* were more robustly upregulated with 18- and 15-fold induction in response to calprotectin treatment and 320- and 227-fold induction after treatment with TPEN (Fig 5 A-C). Expression of *SAK_0514*, ortholog to the cation diffusion facilitator, *czcD,* of *S. pneumoniae,* was also assessed by qRT-PCR to confirm that calprotectin and TPEN were inducing zinc-limiting conditions. As expected, expression of *czcD* was downregulated following exposure to both (Supplemental Fig 1A). To confirm what has been previously described for GBS in zinc-limited chemically defined media (38, 39), we observed that mutants lacking individual zinc-binding proteins exhibited similar calprotectin sensitivity as the WT GBS strain, while growth of a triple Δ*adcA*Δ*adcAII*Δ*lmb* mutant was reduced (Supplemental Fig 1B). This is consistent with previous results that suggest functional redundancy exists between zinc-binding proteins.

**Figure 5.**
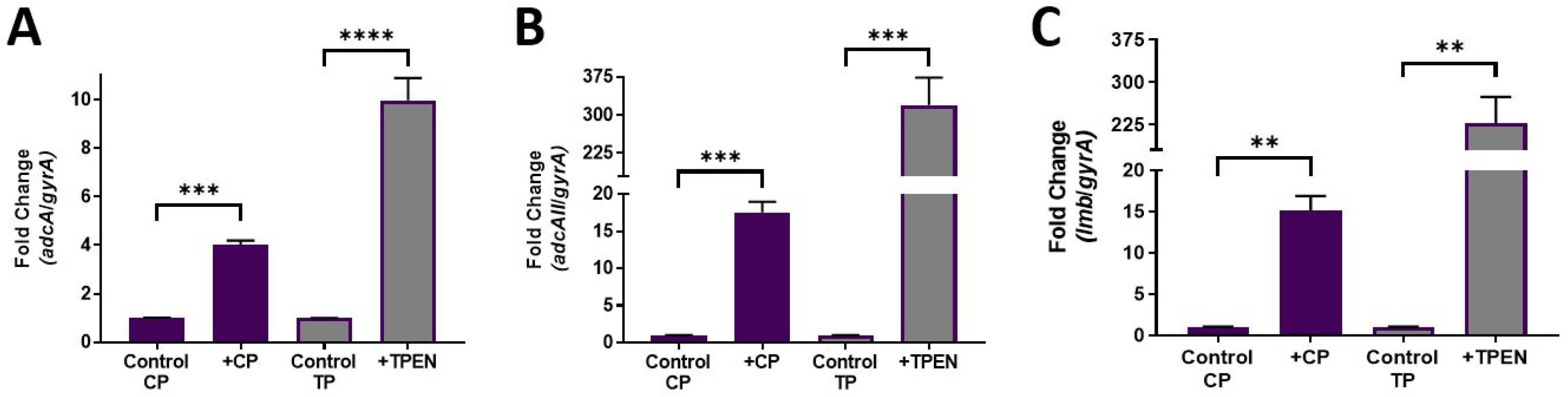
Zinc transport contributes to calprotectin resistance. Quantitative RT-PCR was used to assess expression of **A)** *adcA*, **B)** *adcAII*, and **C)** *lmb* following exposure to 120 μg/mL calprotectin or 25 μM TPEN. Fold change was calculated by ΔΔCT analysis with *gyrA* serving as the internal control. Data are displayed as the average fold change from three independent experiments. Significance was determined by unpaired Student’s t-test, ** *p*<0.01, *** *p*<0.001, **** *p*<0.0001..

### Zinc homeostasis contributes to GBS virulence and meningitis

Our results thus far suggest that GBS utilizes zinc uptake machinery to cope with calprotectin stress *in vitro*; thus, we hypothesized that zinc homeostasis would contribute to GBS virulence. Using a murine model of GBS systemic infection we infected mice (C57BL/6) intravenously with WT or the Δ*adcA*Δ*adcAII*Δ*lmb* mutant strain. Infection with the WT GBS strain resulted in significantly higher mortality than with the isogenic mutant strain (Fig 6A). By 36 hours post-infection, 7/8 WT infected mice succumbed to infection. Conversely, only 3/8 mice challenged with the Δ*adcA*Δ*adcAII*Δ*lmb* strain succumbed to infection by the experimental endpoint of 144 hours (Fig 6A). At the time of death or the experimental endpoint, blood and brain were harvested to determine bacterial load. Despite a similar level of bacterial CFU recovered from blood (Fig. 6B), a significantly higher bacterial burden was observed in brain tissue (Fig 6C), of WT GBS infected animals compared to animals infected with the Δ*adcA*Δ*adcAII*Δ*lmb* mutant strain. Similar results were also observed during infection in another mouse (CD-1) background (Supplemental. Fig 2). We further detected an increase in KC, a neutrophil chemokine, in brain homogenates of mice challenged with WT GBS compared to the Δ*adcA*Δ*adcAII*Δ*lmb* mutant strain (Fig 6D) suggesting a more robust infection and increased inflammation in WT infected mice. To determine the contribution of calprotectin specifically to GBS disease progression, we infected *S100A9*-/- mice with WT and Δ*adcA*Δ*adcAII*Δ*lmb* GBS. We observed that *S100A9*-/-mice were equally susceptible to WT and mutant GBS (Fig 6E) and had similar bacterial loads in the brain and blood (Fig 6F,G) and levels of neutrophilic chemokine, KC in brain tissue (Fig 6H). Interestingly, *S100A9-/-* mice were less susceptible to WT GBS infection compared to WT mice. Taken together these data indicate that GBS zinc homeostasis is required for invasive disease, specifically in the presence of host calprotectin.

**Figure 6.**
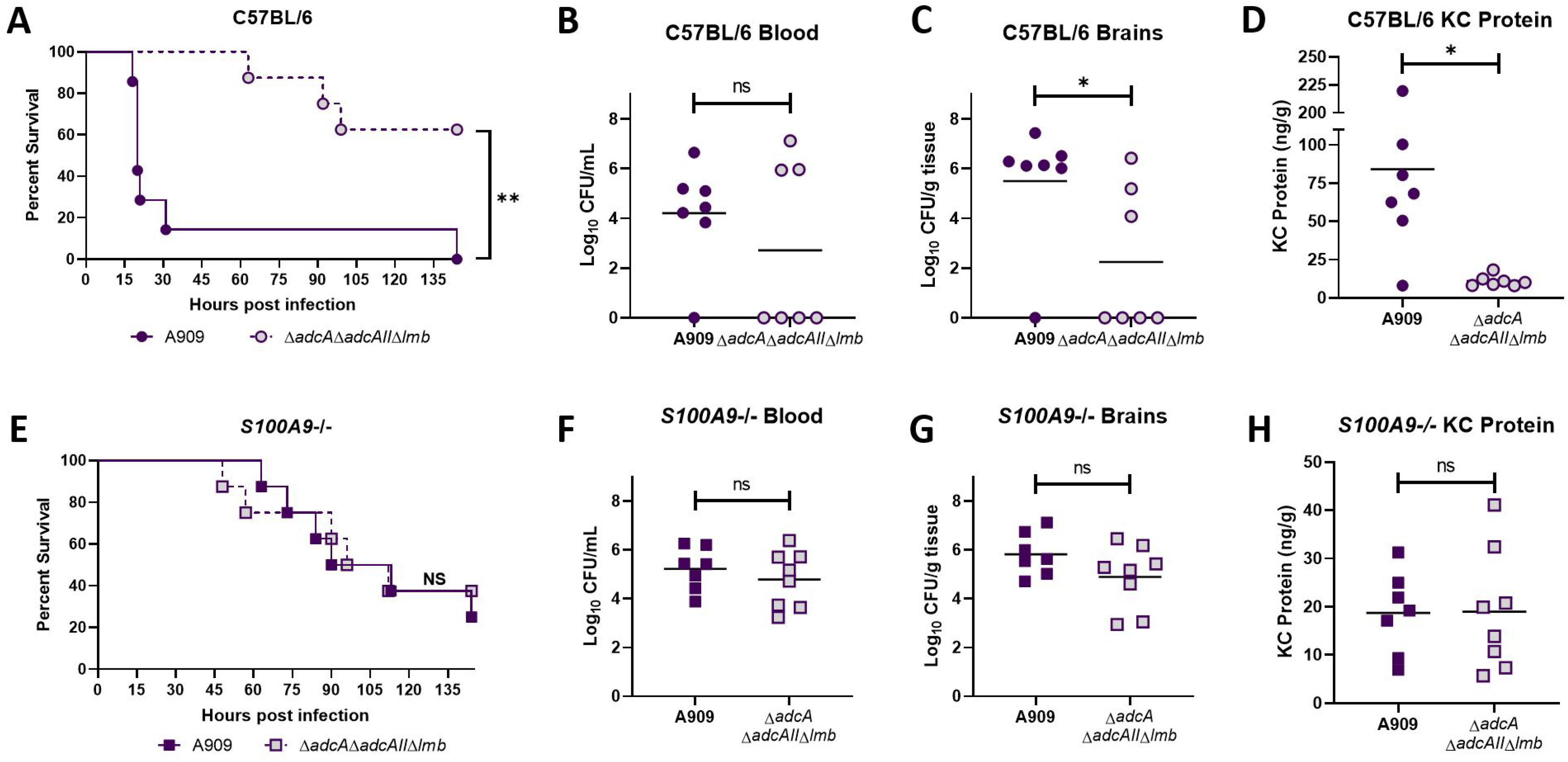
GBS zinc homeostasis contributes to calprotectin survival *in vivo*. Kaplan-Meier plot showing survival of **A)** C57BL/6 or **E)***S100A9*-/- mice infected with 3×10^8^ CFU of WT (solid line) or the Δ*adcA*Δ*adcAII*Δ*lmb* mutant (dotted line). Recovered CFU were quantified from brain tissue homogenates **(B and F)** or blood **(C and G)**. Cytokine abundance was quantified from brain tissue homogenates by ELISA **(D and H)**. Statistical analyses include Log-rank (Mantel-Cox) test for panels A & E and unpaired Student’s t-test for panels B-D and F-H, * *p*<0.05 and ** *p*<0.01.

## Discussion

GBS infections are known to result in increased immune cell influx and inflammation, specifically neutrophilic infiltrate (33, 56). As these are characteristic signs of bacterial meningitis, GBS would encounter high concentrations of granulocyte-derived calprotectin (57) during infection. Calprotectin makes up more than 50% of the neutrophil cytosol, and has been proven to be an effective molecule at starving incoming pathogens of nutrient metal ions (7, 58). However, despite this mechanism employed by the immune system to impede bacterial growth, GBS continues to cause life-threatening illnesses, suggesting that this bacterium possesses machinery to thwart host defenses and permit survival. Here we have examined the global effect of calprotectin stress on GBS fitness using a newly developed *mariner*-GBS transposon mutant library. We identified systems involved in zinc and manganese/iron homeostasis, as well as putative metal-transport systems that have not been previously described in GBS to be important for GBS growth in the presence of calprotectin. Through mutagenesis and functional analyses, we determined that the Adc zinc acquisition system, comprising three zinc-binding proteins promotes survival during calprotectin stress and contributes to systemic infection *in vivo*. Further the loss of calprotectin *in vivo* ablates the requirement of zinc homeostasis for GBS virulence. These data support the growing appreciation for the role of zinc uptake in bacterial pathogenesis and provide new insight into the mechanisms by which GBS resists nutritional immune challenge (Fig 7).

**Figure 7.**
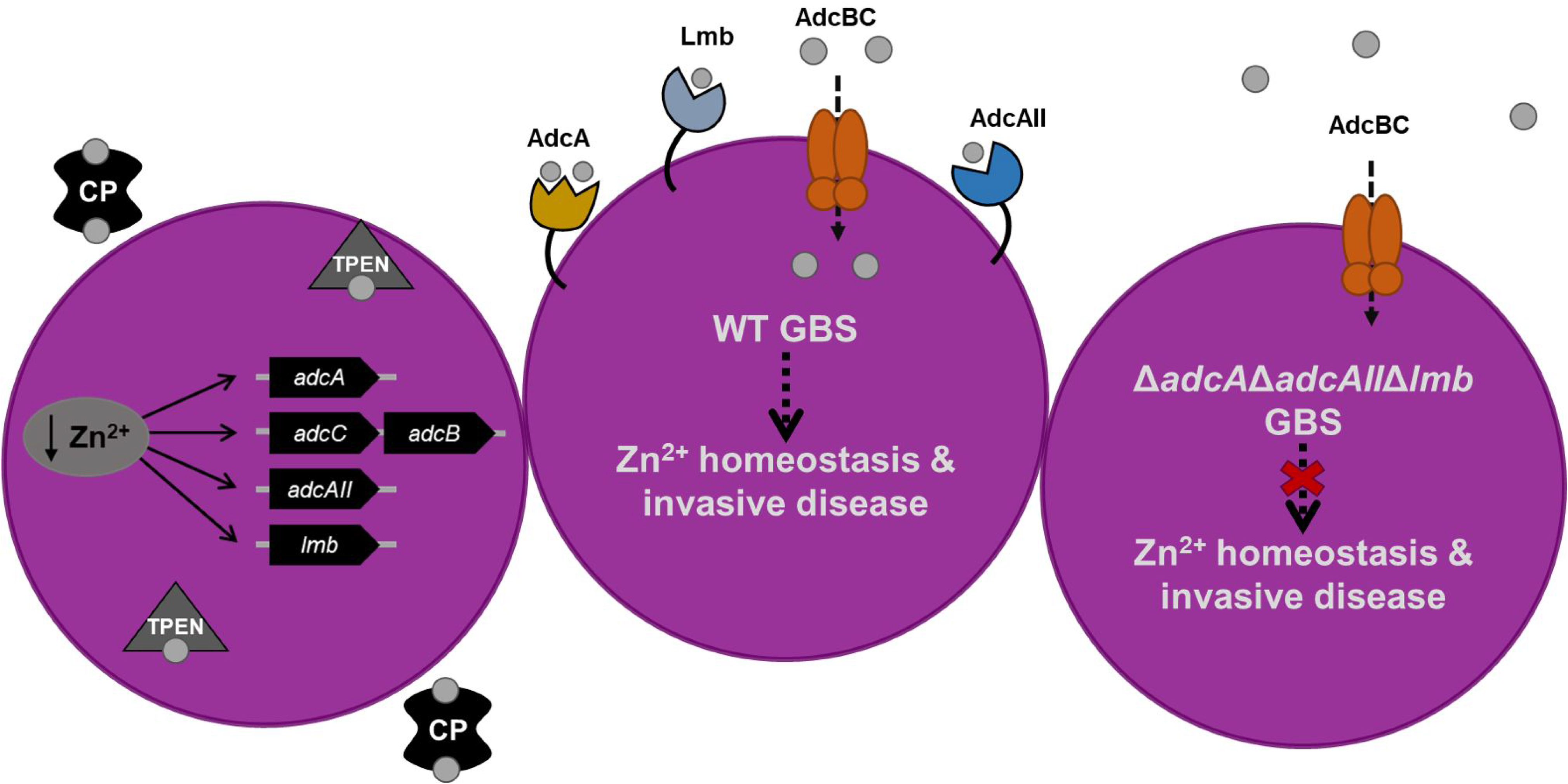
Summary of the GBS zinc-dependent response to calprotectin. GBS senses metal limitation in the presence of calprotectin and induces expression of three zinc-binding proteins to acquire zinc and overcome starvation. GBS that are capable of regulating zinc homeostasis in a zinc-limited environment remain virulent; whereas GBS zinc-transport mutant strains are deficient in their ability to cause invasive disease.

In this study we demonstrate for the first time the global impact of calprotectin-mediated metal chelation on GBS fitness using transposon library screening. We observed that growth of both GBS disease and colonizing clinical isolates was inhibited by physiologically relevant concentrations of recombinant calprotectin. Serotype V isolates were significantly more resistant to chelation than serotype Ia isolates *in vitro*, but the basis for this requires further investigation. As our current understanding of GBS metal homeostasis is limited, we sought to characterize the global effects of calprotectin-mediated metal chelation on GBS fitness, utilizing a newly constructed, *Krmit*-transposon mutant library. This screen identified COG categories of GBS gene function during calprotectin stress, with the most abundant genes of known function involved in inorganic ion transport, amino acid and carbohydrate transport and metabolism, transcription, cell wall biogenesis and defense mechanisms. Some of the most significant underrepresented factors identified were those involved in metal transport including zinc/manganese/iron uptake and efflux. Our data also identified GBS essential genes for growth in rich media, including THY and mRPMI, and our results were consistent with a previous study using transposon sequencing of a different GBS strain, which identified essential genes in tRNA synthesis pathways, glycolysis, and nucleotide metabolism (59). We did however observe some differences. The transcriptional regulator *ccpA* was previously deemed part of the GBS essential genome (59), though in our analysis *ccpA* was only essential in mRPMI. Similarly, the global nutritional regulator *codY*, which is essential for *Streptococcus pneumoniae* growth (60) and nonessential for GBS in rich media (59), was found in our study to be nonessential for growth in THY but essential for GBS growth in mRPMI. Together these data suggest that the essential genome of GBS is largely similar across strains although this is dependent on the growth medium.

In order to survive metal limiting environments, bacterial pathogens possess tightly regulated, high affinity metal uptake systems and many Gram-positive bacteria are known to use the ZnuABC/AdcABC zinc transport systems (61–63). In the case of pathogens such as *S. aureus* and *Bacillus anthracis*, each possesses a single zinc-binding protein, AdcA and ZnuA, respectively (64, 65), and their associated ATP-binding cassette transporter permease and ATPase are encoded by AdcB/ZnuB and AdcC/ZnuC (38, 66).(38, 66) The zinc uptake machinery of streptococcal pathogens *S. pneumoniae* and *S. pyogenes* possess a second zinc-binding lipoprotein, AdcAII. GBS is unique in that it has a third zinc-binding protein, Lmb (67), encoded on a mobile element that is co-transcribed with a second streptococcal histidine triad protein, ShtII. Lmb is thought to have been acquired by horizontal gene transfer and shares homology with Lsp of *Streptococcus pyogenes* (68), but the direct origin remains unknown. Originally annotated as laminin binding proteins, Lmb/Lsp were thought to contribute to adherence, though this interaction has been debated and may be species or strain dependent (42, 69, 70). The GBS zinc uptake machinery is encoded by four distinct operons and is under the regulation of the zinc-dependent AdcR repressor (38). AdcR is involved in maintaining intracellular zinc homeostasis, and has been shown to be important for growth in zinc-limiting conditions (38, 39). This system is also significantly upregulated during GBS murine vaginal colonization and following incubation with human blood (71, 72). In addition to the ABC transporters, bacteria have evolved other mechanisms to maintain zinc homeostasis, examples include the metallophore staphylopine produced by *Staphylococcus aureus* that binds zinc ions and is imported by the CntABCDF machinery (65), and the TdfH transporter of *Neisseria gonorrhoeae* that binds calprotectin directly to hijack and secure zinc ions (73). Though similar systems have not been described in GBS.

Our transposon mutant screen during calprotectin treatment identified loss of function mutations in the zinc-binding protein AdcA as the most significantly underrepresented metal mutation. Mutants in the AdcC subunit of the zinc-dependent ABC-transporter, zinc-binding proteins, AdcA and AdcAII, and the streptococcal histidine triad protein, ShtII were also underrepresented in our screen. Interestingly, mutations in Lmb and Sht did not result in fitness defects in our transposon library screen, suggesting that in a competitive growth environment, loss of either AdcA or AdcAII reduced GBS fitness in the presence of calprotectin. However, in monoculture while loss of all three solute binding proteins sensitized GBS to calprotectin, their individual loss did not. Expression of *adcAII* and *lmb* were both induced to greater extent than *adcA* by both calprotectin and TPEN, and mutations in AdcAII were specifically observed in inhibitory calprotectin treatment. Collectively, these observations could indicate that the GBS zinc importers may uniquely contribute to resisting metal limitation or other aspects of infection, but further investigation is needed.

Additional systems of interest that were underrepresented in our calprotectin transposon library screen were *mtsABC SAK_1554-1556*, *mntH* (*SAK_0871*), *sczA* (*SAK_0515*), *czcD* (*SAK_0514*), and *cadD* (*SAK_2051*). *mtsABC*, encodes the manganese/iron-dependent ABC-transporter and *mtsA* is a component of the core GBS genome and could be a conserved system for survival during calprotectin-mediated stress within the host (51, 74). Additionally, *mntH*, encodes the manganese/iron NRAMP and is known to be important for survival in acidic conditions similar to what GBS would encounter during inflammation or within the phagolysosome (55). Similar to what has been shown for zinc import, the *mtsABC* transporter was shown to be upregulated in human blood and during vaginal colonization (71, 72). In the context of metal ion efflux, we identified mutations in *sczA*, *czcD*, and *cadD*, which all resulted in fitness defects when grown in media containing calprotectin. SczA is a zinc-dependent transcriptional activator of the cation diffusion facilitator protein, CzcD. Together, this system has been shown to contribute to bacterial survival during zinc toxicity, neutrophil and macrophage killing, and contribute to GAS virulence (75–78). To date, SczA has been suggested to be an activator in GBS (75) but the functional roles of SczA and CzcD in metal efflux and GBS survival have not been described. Additionally, the *cadDX* operon in *Streptococcus salivarius* has been shown to have both cadmium and zinc inducible repression (79), thus CadD may function similarly in GBS, but this warrants further investigation. Recently, a new highly virulent GBS sequence type (ST) 485, of the clonal complex 103, has become increasingly common in China, specifically, with the frequency of isolation quickly climbing from 1% to 14% (80). These isolates have evolved from a genetic lineage capable of causing both human and bovine disease, and both the increase in virulence and rapid emergence of these isolates is thought to be due to the acquisition of the *cadDX* operon (80).

An additional strength of our study is the sensitivity of our screen to detect genes involved in overcoming varying degrees of metal starvation. We identified 30 genes that were essential for GBS growth in a subinhibitory concentrations of calprotectin (Supplemental Table 1), representing genes that are necessary for overcoming low level metal sequestration but are nonessential for survival in extreme metal limitation. Genes of importance include those encoding two ribosomal proteins, key enzymes involved in glycolysis, two enzymes involved in folate metabolism, and a cobalt transporter ATPase. We also identified 55 uniquely essential genes for survival in inhibitory levels of calprotectin. These genes represent those that are important for growth when GBS encounters high degrees of starvation or the starvation of multiple metals. Systems of interest in these data were genes encoding four ribosomal proteins, six prophage-related proteins, phosphotransferase systems, and the stress response serine protease, HtrA (Supplemental Table 1). These differential findings are significant as it is becoming increasingly appreciated that metal starvation/intoxication occurs across a gradient and that the maintenance of metal homeostasis is dynamic and requires numerous fine-tuned responses. These data are supported by previous studies that show streptococcal metabolism (81,82) to be dependent on metal ions and that ribosomal proteins serve as reservoirs for intracellular zinc during metal limitation (83).

As the relative contribution of zinc homeostasis to GBS virulence had not been previously characterized, we utilized a murine model of GBS systemic infection to compare WT and Δ*adcA*Δ*adcAII*Δ*lmb* strains. These experiments demonstrated that the Δ*adcA*Δ*adcAII*Δ*lmb* mutant was significantly attenuated compared to WT GBS in two different mouse strains. These data further validate our *in vitro* results and confirm the importance of zinc-uptake machinery to the pathogenesis of GBS infection. To determine the contribution of host calprotectin to the GBS disease process, we utilized a calprotectin knockout mouse strain (*S100A9*-/-) (84, 85). In contrast to the phenotypes observed in WT mice, *S100A9*-/- mice were equally susceptible to GBS WT and Δ*adcA*Δ*adcAII*Δ*lmb* strains (Fig 7A), suggesting that zinc-transport machinery is expendable when the zinc-limiting pressure of calprotectin is absent. Further, we observed that WT mice exhibited increased mortality due to GBS infection compared to *S100A9*-/- mice. In our studies, nearly 90% of WT mice infected with WT GBS succumbed to illness by 48 hours, whereas the *S100A9*-/- mice infected with WT GBS did not reach 50% lethality until 90 hours post-infection. Similar trends were recently observed in *S100A9*-/- mice challenged with *S. pyogenes* (86). Additionally, these data are consistent with previous findings that suggest a role for calprotectin as an immunological alarmin in promoting inflammatory signaling (87–89). Studies to determine the specific role of calprotectin in inflammation during GBS disease progression warrant further investigation.

Here, we report the generation and utilization of a highly saturated GBS *mariner* transposon library to investigate bacterial response to calprotectin-mediated metal chelation. Genome wide screening revealed numerous metabolic pathways and metal transport systems that may contribute to the ability of GBS to overcome calprotectin stress and nutritional immunity. Our results emphasize the importance of zinc-transport to the development of GBS systemic infection, highlighting the significance of zinc homeostasis to disease progression. As zinc uptake machinery are highly conserved across streptococcal pathogens, these present a promising target for the development of novel antimicrobials.

## Materials & Methods

### Bacterial Strains and Growth Conditions

*Streptococcus agalactiae* (GBS) isolates A909 (serotype Ia), CJB111 (serotype V), COH1 (serotype III), a cohort of 25 vaginal colonization isolates (serotypes Ia, Ib, II, III, and V) (47) and mutant strains (A909Δ*adcA*, A909Δ*adcAII*, A909Δ*lmb*, A909Δ*adcA*Δ*adcAII*Δ*lmb*) were cultured in Todd-Hewitt Broth (THB) at 37°C. Mutant strains were constructed as previously described (38, 39). Deletion of these genes did not affect growth in metal sufficient media (38).

### Calprotectin Growth Assays

Briefly, GBS cultures grown overnight for 18h in THB were diluted 1:50 into 100 μL in a 96 well microtiter plate with 38% 3x modified-RPMI (mRPMI) (90), (RPMI powder [Gibco 31800-022], 150 mM HEPES pH 7.4, 1.5% glucose, 3x Basal Medium Eagle (BME) vitamins (Sigma), vitamins, 75 μg/mL guanine, 75 μg/mL uracil, and 75 μg/mL adenine) and 62% calprotectin buffer (20 mM Tris pH 7.5, 100 mM NaCl, 3 mM CaCl_2_), and 0-240 μg/mL recombinant calprotectin (65, 91). At 8 hours post-inoculation, growth was assessed by measuring optical density (OD_600nm_) and plating serial dilutions to quantitate CFU.

### Generation of mariner (Krmit) mutant libraries in GBS for Tn-seq

*In vivo mariner* transposition for random mutagenesis in GBS was accomplished using the pKrmit system originally developed for GAS (48), as previously described (48, 92). Briefly, GBS CJB111 cells were transformed by electroporation with 300 μg of pKrmit and outgrown in THY broth at 30°C (permissive temperature for pKrmit replication) for 4 h. Transformants were then selected by plating on THY agar containing kanamycin, 300μg/mL, and spectinomycin 100μg/mL at 30°C for 48 h. The presence of intact pKrmit was phenotypically tested as previously described (92) and proper GBS transformants were stored at −80°C. For *Krmit* transposition, an individual pKrmit-containing GBS freezer stock was used to inoculate 250 ml of THY broth containing Km and incubated overnight at 37°C (T_0_) (non-permissive temperature for pKrmit replication). The quality of *Krmit* transposition was tested as previously described (92, 93): the complexity or randomness of *Krmit* mutant libraries was assessed by amplifying the insertion sites of random mutants by AP-PCR, Sanger DNA sequencing and mapping onto the appropriate GBS genome. Percent randomness was determined by defining a ratio of unique insertions (by AP-PCR sequencing) among a tested population.

### Transposon library screening

GBS pooled *Krmit* library was grown overnight in THB with kanamycin 300 μg/mL. Overnight culture was back diluted into 38% 3x mRPMI with 62% calprotectin buffer with 0, 60, or 480 μg/mL purified calprotectin. Samples were incubated at 37°C for 8 hours. Following incubation, samples were centrifuged and resuspended in 200 μL PBS. Total volume was spread onto THA plates and incubated overnight at 37°C. Bacterial growth from each treatment condition was collected and pooled into a three samples and genomic DNA was extracted using Zymobiomics DNA miniprep kit (Zymo Research).

### Transposon library sequencing

Library preparation and sequencing was performed as previously described (49) by the Microarray and Genomics Core at the University of Colorado Anschutz Medical Campus. Briefly, genomic DNA was sheared to approximately 340 bp fragments and processed through the Ovation Ultralow V2 DNA-Seq library preparation kit (Tecan) and 9 ng of each library was used as template to enrich by PCR (16 cycles) for the transposon insertions using *Krmit*-specific (TCGTCGGCAGCGTCAGATGTGTATAAGAGACAGCCGGGGACTTATCA**T**CCAACC) and Illumina P7 primers. The enriched PCR products were diluted 1:100, and 20 μL used as template for an indexing PCR (9 cycles) using the TruSeq P5 indexing and P7 primers. Sequencing was performed using Illumina NovaSeq 6000 in 150 base paired-end format.

### Bioinformatic Analyses of Tn-seq

Two annotated *Streptococcus agalactiae* genomes were used as the reference for these analyses: CJB111 (AAJQ01000001) and A909 (NC_007432). Provided annotation is based on the A909 genome as the annotated CJB111 genome has significantly lower coverage. Italicized directory names (ending in /) refer to directories within the git repository for this project, github.com/abelew/sagalacticae_2019. Many of the post-processing is handled by the hpgltools R package (94), functions used are italicized and suffixed with (). Approximately 2.5 million reads of each raw library were queried for quality with Fastqc (95) before removing the *mariner* ITR leading sequences with cutadapt (96). These libraries were aligned against the reference genomes with bowtie (97, 98) using options to allow one mismatch (-v 1) and randomly assign multi-matched reads to one of the possible matching positions (-M 1). The resulting alignments were converted to sorted/compressed binary alignments (99) and counted (100) against the reference genome CDS and intergenic regions. The essentiality software package (50) provides an opportunity to query statistically significant stretches of Tas that have no observed insertions to further inform its metric of essentiality. The insertion data was therefore converted into its expected format and passed to the version 1.21 of the implementation python script. The resulting table provided a count of the number of insertions observed in each ORF, the number of observed Tas, the maximum length of non-observed sequence, the nucleotide span of this region, a call on whether each ORF is essential, and the posterior probability for each call. The default options were used except multiple runs were performed with the minimum hit parameter set to: 1,2, 4, 8, 16, 32. These operations were performed via CYOA. In a separate invocation, the three replicates for the control, low concentration, and high concentration samples were concatenated into a single sample and essentiality was run on the combined samples. The libraries were quantified with respect to relative coverage, similarity, and saturation with respect to available TA insertion points. These tasks were performed using the hpgltools and the input text/wig files for the essentiality package. Thus, the essentiality input files were read into the R function *plot_saturation()* and used to visualize the saturation of each library. This was done by taking the log2(hits + 1) for each position and plotting them as a set of histograms. Comparison and normalization of control (input) and experimental (output) libraries was performed similar to the essentials software package (101, 102), but using a combination of voom/limma, EdgeR, DESeq2, EBSeq, and a statistically uninformed basic analysis instead of EdgeR. Pairwise Euclidean distances, Spearman correlation coefficients, and principle component analyses were then used to visualize the similarities/differences between normalized libraries. Clustering of Orthologous Groups of proteins (COGs) were assigned using EggNOG 5.0.0 (103) and Venn diagrams were calculated and plotted using BxToolBox (www.bioinforx.com).

### Quantitative Reverse Transcriptase PCR (qRT-PCR) and ELISA

GBS were grown to mid-logarithmic phase (OD_600nm_ 0.4) in mRPMI media and incubated with calprotectin 120 μg/mL for 1 hour at 37°C or TPEN 25 μm for 15 minutes at 37°C. Following incubation, bacteria were centrifuged at 5000 x g for 5 minutes, total RNA was extracted (Macherey-Nagel) and cDNA was synthesized (Quanta Biosciences) per manufacturers’ instructions. Primers used in this study are outlined in Table S2. KC from mouse brain homogenates was quantitated by enzyme-linked immunosorbent assay per manufacturer’s instructions (R&D Systems).

### Ortholog Clustering

To evaluate if metal transport machinery involved in survival during calprotectin stress were orthologous to characterized systems in the closely related *Streptococcus pneumoniae* TIGR4 (GCF_000006885.1) and *S. agalactiae* A909 (GCF_000012705.1) was used as input for the program OrthoFinder v. 2.2.6 (104). Domain architecture of proteins in each orthogroup collected were evaluated using InterProScan v. 5.27-66.0 (105). Predicted operons in GBS were determined using the Database of prOkaryotic OpeRons (DOOR^2^) (106–108).

### Mouse model of GBS systemic infection

All animal experiments were conducted under the approval of the Institutional Animal Care and Use Committee (#00316) at the University of Colorado Anschutz Medical Campus and performed using accepted veterinary standards. We utilized a mouse model of systemic infection as previously described (26, 34, 35), where female eight-week old C57BL/6, C57BL/6 *S100A9*-/-, and CD1 mice were injected intravenously with 3×10^8^ CFU of wildtype A909 or the A909Δ*adcA*Δ*adcAII*Δ*lmb* mutant. Mice were euthanized and blood, lung, and brain tissues were collected. Tissue homogenates and blood were plated on THA to quantify GBS CFU burden.

### Statistical analyses

Significance during calprotectin growth experiments was determined by Kruskal-Wallis test with Dunn’s multiple comparisons post-test with treated samples compared to untreated controls. Normality was confirmed for clinical isolate data by Shapiro-Wilk test and significance was determined by One-way ANOVA with Tukey’s multiple comparisons post-test. Calprotectin growth assays using GBS WT and mutant strains was analyzed by Kruskal-Wallis test with Dunnett’s multiple comparisons test. Statistical differences in murine experiments were determined by Log-rank Mantel-Cox test for survival and by unpaired Student’s T-test. Statistical significance was accepted when *p* < α, with α = 0.05.

### Data availability

Sequencing reads from the transposon sequencing analyses are available in the NCBI Sequence Read Archive (SRA) under the accession number (pending).

## Acknowledgements

We want to thank the University of Colorado Anschutz Medical Campus Genomics and Microarray Core and Dr. Alexander Tice at Mississippi State University for assistance with sequencing and data analysis. This work was supported by NIH 5T32AI007405-28 to B.L.S, NIH/NIAID R21 AI134078 and R01 AI047928 to KSM, and NIH/NINDS R01 NS116716 to K.S.D.

## Figure Legends

**Supplemental Figure 1. A)** Quantitative PCR was used to assess expression of *czcD* (*SAK_0514*) following exposure to 120 μg/mL calprotectin or 25 μM TPEN. Fold change was calculated by ΔΔCT analysis with *gyrA* serving as the internal control. Data are displayed as the average fold change from three independent experiments. Statistical significance for panel A was determined by unpaired Student’s t-test, ** *p*<0.01. **B)** Growth of WT and GBS zinc-transport mutant strains was measured by optical density (OD_600nm_) following an 8-hour incubation with recombinant calprotectin (120 μg/mL). Experiments were performed in technical triplicates of *n*=3 and data were averaged together from three independent experiments. Significance was determined by Kruskal-Wallis test with Dunn’s multiple comparisons post-test, * *p*<0.05.

**Supplemental Figure 2. A)** Kaplan-Meier plot showing survival of CD-1 mice infected with 3×10^8^ CFU of WT (solid line) or the Δ*adcA*Δ*adcAII*Δ*lmb* mutant (dotted line). Recovered CFU were quantified from brain tissue homogenates **(B)** or blood **(C)**. Statistical analyses include Log-rank (Mantel-Cox) test for panel A and Unpaired t-test for panels B-C, *** *p*<0.001.

---

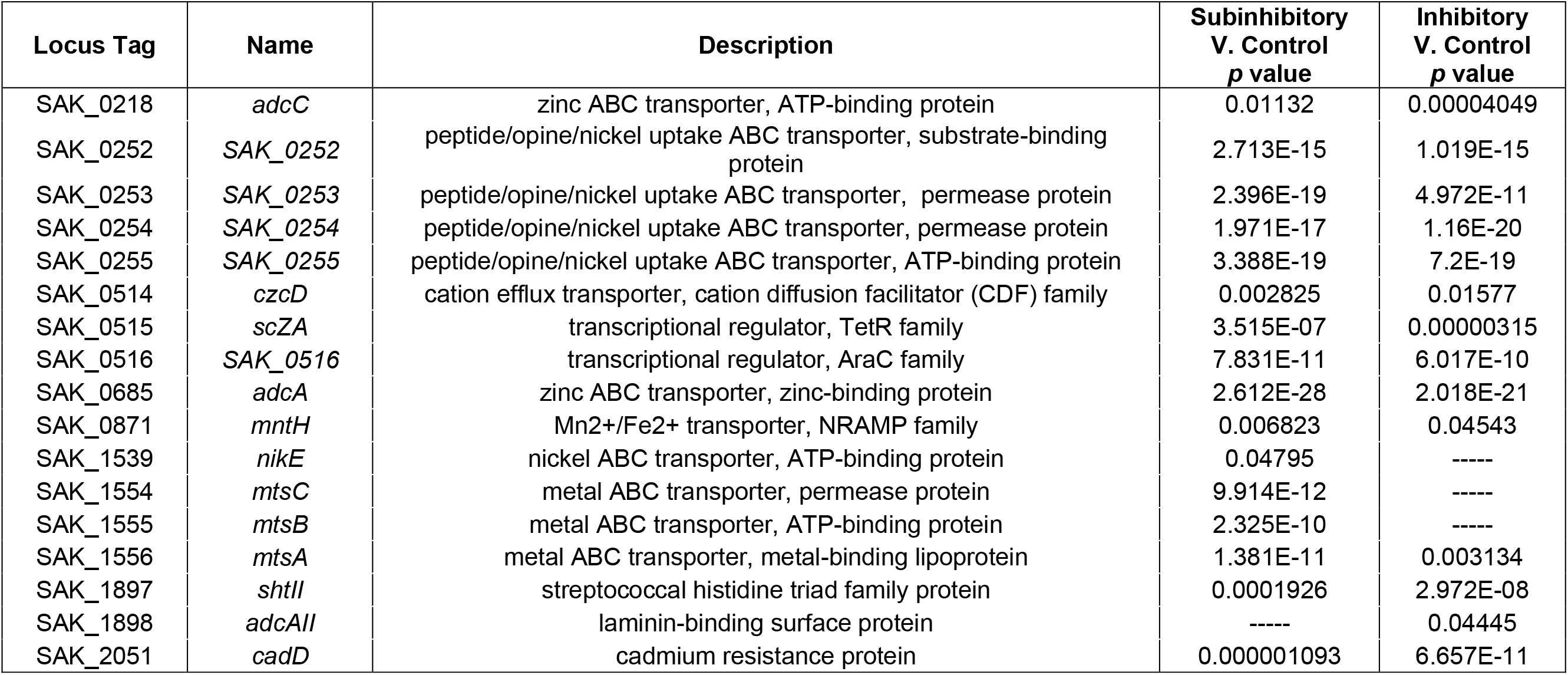

